# Assessment of the performance of different hidden Markov models for imputation in animal breeding

**DOI:** 10.1101/227157

**Authors:** Andrew Whalen, Gregor Gorjanc, Roger Ros-Freixedes, John M Hickey

## Abstract

In this paper we review the performance of various hidden Markov model-based imputation methods in animal breeding populations. Traditionally, heuristic-based imputation methods have been used for imputation in large animal populations due to their computational efficiency, scalability, and accuracy. However, recent advances in the area of human genetics have increased the ability of probabilistic hidden Markov model methods to perform accurate phasing and imputation in large populations. These advances may enable these methods to be useful for routine use in large animal populations. To test this, we evaluate here the accuracy and computational cost of several methods in a series of simulated populations and a real animal population. We first tested single-step (diploid) imputation, which performs both phasing and imputation. Then we tested pre-phasing followed by haploid imputation. We tested four diploid imputation methods (fastPHASE, Beagle v4.0, IMPUTE2, and MaCH), three phasing methods, (SHAPEIT2, HAPI-UR, and Eagle2), and three haploid imputation methods (IMPUTE2, Beagle v4.1, and minimac3). We found that performing pre-phasing and haploid imputation was faster and more accurate than diploid imputation. In particular, we found that pre-phasing with Eagle2 or HAPI-UR and imputing with minimac3 or IMPUTE2 gave the highest accuracies in both simulated and real data.

## Introduction

In this paper we review and analyse the use of hidden Markov model (HMM) based imputation methods for animal breeding populations. Genotype imputation is a key aspect of many modern animal breeding programs and allows genetic information to be obtained on a large number of animals at a low cost. When imputation is applied to a breeding program, a small subset of individuals (e.g., sires) are genotyped at high density, and the remaining animals are genotyped at a lower density. Statistical regularities between shared chromosomal segments are used to fill in the untyped loci. Modern imputation methods fill in missing genotypes at a very high accuracy (e.g., Hickey et al., 2012; Sargolzaei et al., 2011), increasing the number of animals that can be genotyped for a fixed budget. The larger pool of genotyped animals increases the accuracy of genetic predictions on all animals (Daetwyler et al., 2008) and offers the potential to increase selection intensity.

Traditionally, heuristic imputation methods have dominated animal breeding (Hickey et al., 2012; Sargolzaei et al., 2011; VanRaden et al., 2013). These heuristic methods use large chromosome segments shared between closely related animals to rapidly and accurately impute untyped or otherwise missing loci. In contrast, imputation methods used in human genetics have largely been based on the probabilistic HMM framework of Li and Stephens (2003). These probabilistic methods tend to have higher accuracy than heuristic methods in datasets where individuals are not closely related. However, these methods have come at too high of a computational cost for routine imputation in animal populations.

In the last few years, the speed of HMM methods has improved. They have been used to impute hundreds of thousands of individuals to hundreds of thousands of loci in reasonable computational time (Browning and Browning, 2016; Loh et al., 2016a). These improvements have been driven by the widespread availability of large haplotype reference panels, and the emergence of a two-step imputation pipeline where observed genotypes are first phased and then untyped loci are imputed based on their phased haplotypes (Spiliopoulou et al., 2017). The improved scaling of HMMs may allow for their routine use in large animal breeding populations. However, given the lack of appropriate public domain haplotype reference panels for many animal populations, smaller population sizes, and sparser marker density, it is not clear that the advances in HMMs will be realized for animal imputation. Furthermore, there are a number of competing HMM imputation methods and it is not clear which is most suited for routine use in animal breeding.

In this paper we provide a high-level review of several imputation methods and study their performance on simulated and real data. We grouped comparisons based on single-step (diploid) imputation methods and a two-step combination of pre-phasing and haploid imputation methods. Specifically, for diploid imputation we test fastPHASE (Scheet and Stephens, 2006), Beagle v4.0 (Browning and Browning, 2007), IMPUTE2 (Howie et al., 2009), and MaCH (Li et al., 2010). For pre-phasing we test SHAPEIT2 (Delaneau et al., 2012), HAPI-UR (Williams et al., 2012), and Eagle2 (Loh et al., 2016b), followed by haploid imputation with IMPUTE2 (Howie et al., 2009), Beagle v4.1 (Browning and Browning, 2016), or minimac3 (Das et al., 2016). We first review these methods and then evaluate the performance of these methods on simulated and real data.

## Hidden Markov Models

All of the methods considered are based on Li and Stephens’ (2003) HMM framework. Under this framework an individual’s genotype is considered to be a mosaic of haplotypes from a haplotype reference panel *H=(h_1_…h_K_}*. The methods calculate the probability that the individual has the pair of haplotypes, *h_j_* and *h_k_* at a locus *i* given the observed genotype (*g_i_*), *p(h_ij_, h_ik_*/*g*_i_). To account for linkage between adjacent loci, the methods evaluate the probability of a haplotype based on its fit to the observed genotypes at the loci and its similarity to the haplotypes inferred at nearby loci:

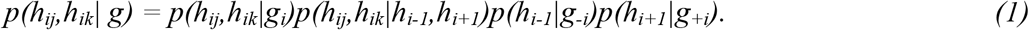

The term *p(h_ij_,h_ik_*/*g_i_*) measures the fit between the pair of haplotypes and the observed genotype at a locus. The term *p(h_ij_,h_ik_*/*h_i-1_,h_i+1_)* captures transitions between haplotypes given the haplotypes at neighbouring loci. The terms *p(h_i-1_*/*g_-i_)* and *p(h_i+1_*/*g_+i_)* measure the fit between haplotypes and observed genotypes at the remaining loci. These probabilities can be calculated using the standard forward-backward algorithm (Rabiner, 1989).

Traditionally, methods that rely on the Li and Stephens framework scale linearly with both the number of individuals and the number of loci and quadratically with the number of reference haplotypes. The quadratic scaling is due to phase uncertainty at heterozygous loci, requiring the methods to model haplotypes assigned on both chromosomes simultaneously. The quadratic scaling quickly leads to intractable computational costs even for small reference panels, but can be avoided if the low-density individuals are pre-phased, which allows haplotypes to be considered independently. Haploid imputation, imputation with pre-phased haplotypes, therefore scales linearly with the number of individuals, number of loci, and number of reference haplotypes.

In this paper we consider two classes of HMMs. In the first class, diploid imputation methods perform phasing and imputation simultaneously, resulting in quadratic scaling with the reference panel size. To mitigate this issue, each of the evaluated methods, fastPHASE, Beagle v4.0, IMPUTE2, and MACH, employ their own strategy to reduce the effective number of reference haplotypes while maintaining high accuracy. In contrast, two-step imputation methods treat phasing and imputation as separate problems. Individuals are first phased and then imputed using a haploid HMM which scales linearly with the number of reference haplotypes. Phasing methods may have either quadratic, super-linear, or linear dependence on the number of reference haplotypes. A number of tricks are deployed to increase phasing speed and accuracy that would not be applicable if the phasing methods also needed to handle genotype uncertainty at untyped loci.

Intuitively, we might expect that the diploid imputation methods will have higher accuracy (at a higher computational cost) than separately performing phasing and imputation because they automatically handle phase uncertainty. This is not necessarily the case if most errors in imputation stem from the inability to find appropriate reference haplotypes that would explain observed genotypes. By performing pre-phasing and then imputation, it may be possible to consider a much larger number of reference haplotypes and thereby increase accuracy by finding a more appropriate set of reference haplotypes which offset accuracy losses due to phasing errors.

Below we review methods for diploid imputation, haploid imputation, and phasing.

### Diploid imputation

All four diploid imputation methods utilize a haplotype state-space reduction technique to alleviate the impact of modelling a large number of haplotype reference panels. IMPUTE2 and MaCH use subsampling, where the haplotypes considered in each iteration are a sample of the total haplotype pool. fastPHASE and Beagle v4.0 use haplotype clustering, where the overall number of haplotypes is collapsed into a smaller number of “ancestral” haplotypes.

In the case of IMPUTE2 and MaCH, each method is run over a series of iterations, and at each iteration a subset of the haplotype reference panel is used to phase and impute individual’s genotypes. In MaCH, the subset is selected randomly. In IMPUTE2, the subset is selected to be made up of haplotypes that are “nearby” the currently estimated haplotype for the individual. If these methods are run without an external reference panel, a reference panel is built up from the current phasing of high-density individuals. At each iteration, a new subset of the reference panel is selected for each individual, individuals are imputed and phased based on that subuset, and then a reference panel is re-computed from the currently inferred haplotypes. The methods are run for a small number of iterations (e.g., 20) and the imputation results are averaged across iterations. There is a potential danger in applying these methods in populations of many closely related individuals, due to the potential for feedback between the phasing of closely related relatives (Nettelblad, 2013).

In contrast, in fastPHASE and Beagle v4.0 individuals are imputed based on a set of estimated “ancestral” haplotypes. In fastPHASE, an expectation-maximisation (EM) algorithm is used to infer a small number of ancestral haplotypes from the data (e.g., 30) and then iterates between estimating the haplotypes of each individual as a mosaic of ancestral haplotypes, and estimating the ancestral haplotypes based on the haplotype assignments of each individual. Beagle v4.0 uses a similar approach as fastPHASE, but instead of using a fixed number of ancestral haplotypes, it infers the number of ancestral haplotypes at each marker and models the transition between ancestral states at adjacent markers in the form of a directed acyclic graph.

### Haploid imputation

In contrast to the four diploid methods, haploid methods do not need to use a state-space reduction technique to handle moderate numbers of haplotypes, because they consider each phased chromosome independently and scale linearly with the number of haplotypes in the reference panels. However, with the recent focus of imputing large bio-bank size human populations (over 100,000 individuals) to whole genome sequence level data, many of the current haploid methods utilize techniques to reduce the computational burden when analyzing large numbers of individuals at a large number of markers.

The haploid HMM used by Impute2 is a straightforward extension of the diploid method implemented in the same program. It uses a subset of haplotypes (based on their similarity to the individual’s current phasing) to impute individuals. Minimac3 uses a similar technique, but instead of subsetting the reference panel it uses a loss-less haplotype compression technique that combines haplotypes that are identical in a region and updates the likelihood of those haplotypes simultaneously. This update is particularly useful for whole genome sequence data where there may be limited haplotype variation over long windows. Beagle v4.1 moves away from the graph-based haplotype model in Beagle v4.0 and uses a more traditional Li and Stephens model. To reduce computational burden, Beagle v4.1 aggregates adjacent loci together into strings and performs updates based on strings instead of individual markers. In addition it only updates the haplotype probabilities at genotyped loci and linearly interpolates the haplotype probabilities at untyped loci.

### Pre-phasing methods

Just as with diploid imputation, HMM-based phasing methods naively scale quadratically with the number of haplotypes in the reference panel. However, this quadratic scaling can be avoided by a state-space reduction technique of splitting the chromosomes into small windows, and assuming that linkage information decays quickly across the window boundaries. Both SHAPEIT2 and HAPI-UR utilize a window-based approach, whereas Eagle2 manages the quadratic dependence by performing a limited beam search through the haplotype space.

SHAPEIT2 operates by splitting the chromosome into small haplotype windows, each containing three heterozygous loci. For each window, there are 2^3^=8 possible ways to phase it, and there are 2^6^=64 possible transitions between windows. SHAPEIT2 evaluates the probability of each of the 8 possible haplotypes and 64 transitions based on a haplotype reference panel, and then phases individuals by sampling haplotypes based on their posterior probabilities. The probability of a haplotype in a given window, and transition between windows can be evaluated in a time that scales linearly with the number of reference haplotypes. As in IMPUTE2, SHAPEIT2 subsets the haplotype reference panel by selecting haplotypes that are nearby the current haplotypes of the individual.

The window splitting approach may lead to reduced accuracy in animal breeding populations, where individuals are expected to share long chromosome segments. In SHAPEIT2 only the between-window transmission probabilities are modeled, and not the probabilities of the underlying reference haplotypes. This means that haplotype assignment information from a given window is only used to update the next window and is ignored for further windows. This approach limits the amount of long range haplotype information (covering more than 3 heterozygous loci) that can be exploited. One solution to this is to increase the size of the windows.

HAPI-UR takes a similar approach to SHAPEIT2 in reducing the large state-space, but uses a series of growing windows which allow it to exploit longer shared chromosomal segments. In order to process large windows, HAPI-UR takes advantage of a number of computational tricks to drastically reduce computation time. Unlike most methods that assume a small error rate for observed genotypes (to cover genotyping errors, errors in the reference panel, and mutations from the ancestral state), HAPI-UR sets the probability of all reference haplotypes that disagree with the observed haplotype to 0. This allows the evaluation of which haplotypes fit an individual’s chromosome to be re-formulated as a bit-wise set-intersection operation. In addition to this, HAPI-UR uses a structured representation of the reference haplotypes that allows for fast lookups of matching haplotypes, and for each individual creates individual specific diploid HMM, which ignores all haplotypes that disagree with homozygote sites. Instead of using a fixed window size, HAPI-UR uses dynamic windows which start small (4 markers) and grows to a user specified maximum (e.g. 64 markers) allowing the method to capture longer chromosome segments.

Eagle2 takes a different approach to phasing individuals by not using a window-based haplotype representation. Instead Eagle2 uses a highly efficient reference haplotype storage method based on the positional Burrows-Wheeler Transform (Durbin, 2014) to allow for looking up consistent haplotype pairs in constant time. Instead of employing a full HMM to evaluate all possible haplotypes, Eagle2 employs a beam search to search through only the most promising paths through the space of all possible haplotype pairs. At each heterozygous locus, these paths branch into two possible sub-paths based on the two phasing options. Low probability paths are pruned or merged to keep the overall number of paths small. To decrease the impact that errors in one part of the genome have on subsequent paths, haplotypes are called after 20 markers allowing for the back-propagation of relevant genetic information while decreasing the potential impact of genotyping errors. Absence of approximate window-based haplotype representation makes Eagle2 particularly appealing for animal populations, where a large number of close relatives share long chromosome segments.

## Materials and Methods

We evaluated the performance of the four diploid imputation methods, fastPHASE, Beagle v4.0, IMPUTE2, and MaCH and the three phasing methods, SHAPEIT2, HAPI-UR, and Eagle2 followed by three haploid imputation methods, IMPUTE2, Beagle v4.1, and minimac3 on a series of simulated datasets and a real dataset.

The simulated dataset modelled a cattle population. The population consisted of 5 generations of 2,000 animals, genotyped on a single chromosome. Each generation was produced by selecting 100 sires from the previous generation based on their true breeding values and randomly mating them with 1,000 dams. The initial set of haplotypes was sampled using a Markovian Coalescent Simulator (Chen et al., 2009) assuming a single 100-cM long chromosome simulated using a per site mutation rate of 2.5□ × □10^−8^, and an effective population size (Ne) that changed over time. Based on estimates for the Holstein cattle population (Villa-Angulo et al., 2009), the Ne was set to 100 in the final generation of simulation and to 1256, 4350, and 43 500 at 1000, 10 000, and 100 000 generations ago, with linear changes in between. The simulation of breeding values and progeny’s haplotypes were performed using AlphaSim (Faux et al., 2016).

In the baseline scenario, a single chromosome was genotyped either with a high-density array of 1,000 SNP (allele frequency greater than 0.01) or with a low-density array of 200 SNP, evenly spaced across the high-density array. All of the sires and 100 dams were genotyped at high density. The remaining animals were genotyped at low density. To test the robustness of each method we independently modified the baseline scenario by varying:

- the number of SNP in the low-density array from 5 to 400,
- the number of individuals in the population from 200 to 10,000, and
- the number of genotyped dams from 0 to 500.
- We also considered the case when the first two generations were genotyped on a different high-density array from the next two generations, with either 25, 50, or 75% of SNP overlapping between the two high-density arrays.

To compare the methods on a real data set, we performed imputation on 56,607 individuals from a commercial pig breeding program. These animals were genotyped either with a high-density array of 60,000 SNP or 80,000 SNP or a low-density array of 15,000 SNP. To estimate imputation accuracy, we selected 500 high-density animals (typed at 60,000 SNPs) and masked them to mimic the pattern of missingness found in the SNP of 500 low-density animals. We restricted imputation to chromosome 1.

Accuracy was measured with the correlation between animals’ imputed genotypes and their true genotypes for each animal separately and averaged over all animals. We did not assess phase accuracy independent of the resulting imputation accuracy.

For the simulated datasets, each method was given 8GB of memory and 24 hours to run. Jobs were terminated if they exceeded the runtime or the memory requirements. Unless otherwise specified, we used the default parameters for each simulation. We tested IMPUTE2 using either the default 10-cM windows or the entire chromosome and found that imputing the entire chromosome increased accuracy at the cost of additional computational time. We used 5-cM windows with an overlap of 1 cM for Beagle v4.0 and Beagle v4.1. The real dataset was imputed with only the two-step imputation methods given their high accuracy and low runtimes.

In all cases, the high-density individuals and low-density individuals were phased separately. For the case of multiple high-density arrays, we used the “merge_ref_panels” option in IMPUTE2 and phased both high-density arrays separately. Because neither minimac3 or Beagle v4.1 accept multiple high-density arrays, we phased the high-density individuals together and let the phasing method fill in the missing genotypes for high-density individuals.

## Results

### Accuracy

The performance of diploid imputation methods is given in Figure 1. Among the diploid imputation methods, MaCH performs well in most settings. Its accuracy depends slightly on the number of high-density dams, the number of low-density SNPs, and the overlap between high-density arrays. The performance of fastPHASE was similar to that of MaCH, but performed better when there were a small number of high-density animals or small overlap between high-density arrays. IMPUTE2 had similar accuracy to MaCH, but performed worse when given a small number of high-density dams, or a small number of individuals, and performed better than MaCH when a large number of high-density dams were given. Beagle v4.0 performed similarly to IMPUTE2, but was less affected by the number of high-density dams and number of individuals.

**Figure 1.**
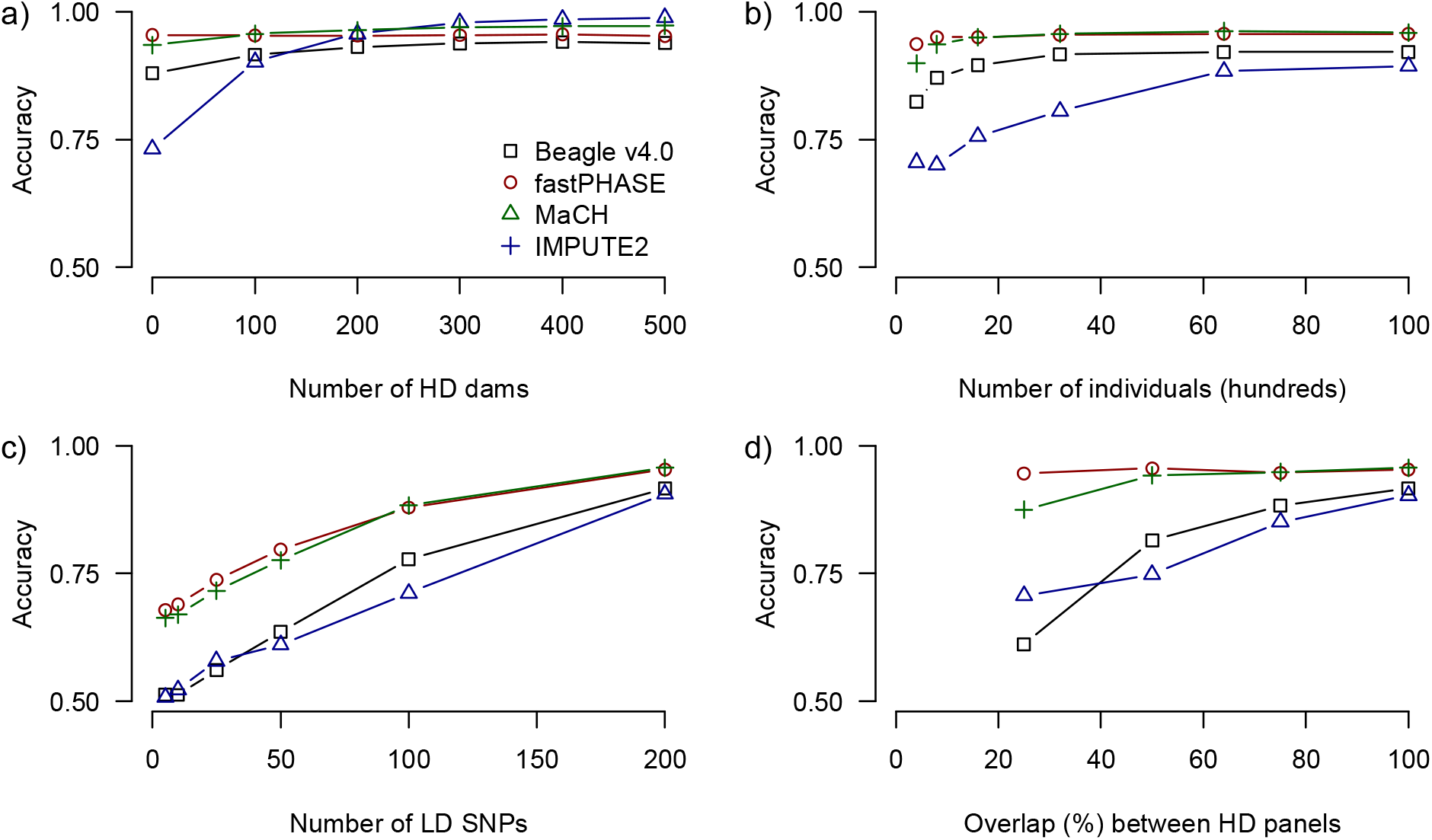
*Performance of each diploid HMM algorithm for each set of simulations. Unless otherwise noted there were 1000 high-density SNPs, 200 low-density SNPs, 100 dams genotyped at high-density and complete overlap between the high-density arrays of generations 1 and 2 and those of 3 and 4. We varied (a) the number of dams genotyped at high-density, (b) the number of individuals in the population, (c) the number of SNPs in the low-density array and (d) the amount of overlap between the high-density array for generations 1 and 2 and those of 3 and 4*.

The performance of pre-phasing and haploid imputation methods is given in Figure 2. Among these methods, we found that the combination of Eagle2 and IMPUTE2 gave the highest imputation accuracy. Eagle2 led to the highest downstream imputation accuracy regardless of the imputation method, and led to higher accuracies than any of the diploid imputation methods. SHAPEIT2 led to similar but slightly lower performance than Eagle2. HAPI-UR led to the lowest overall performance. Of the tested haploid imputation methods we found only a small difference between IMPUTE2 and Minimac3, but found that Beaglev4.1 had poor imputation accuracy in all tested scenarios. We re-ran Beagle v4.1 with different-sized windows but did not see a noticeable increase in accuracy. There was no interaction between the choice of phasing method and the choice of imputation method for the overall imputation accuracy with the exception of when multiple high-density arrays were used. In this case the combination of HAPI-UR and minimac3 outperformed the combination of Eagle2 and minimac3.

**Figure 2:**
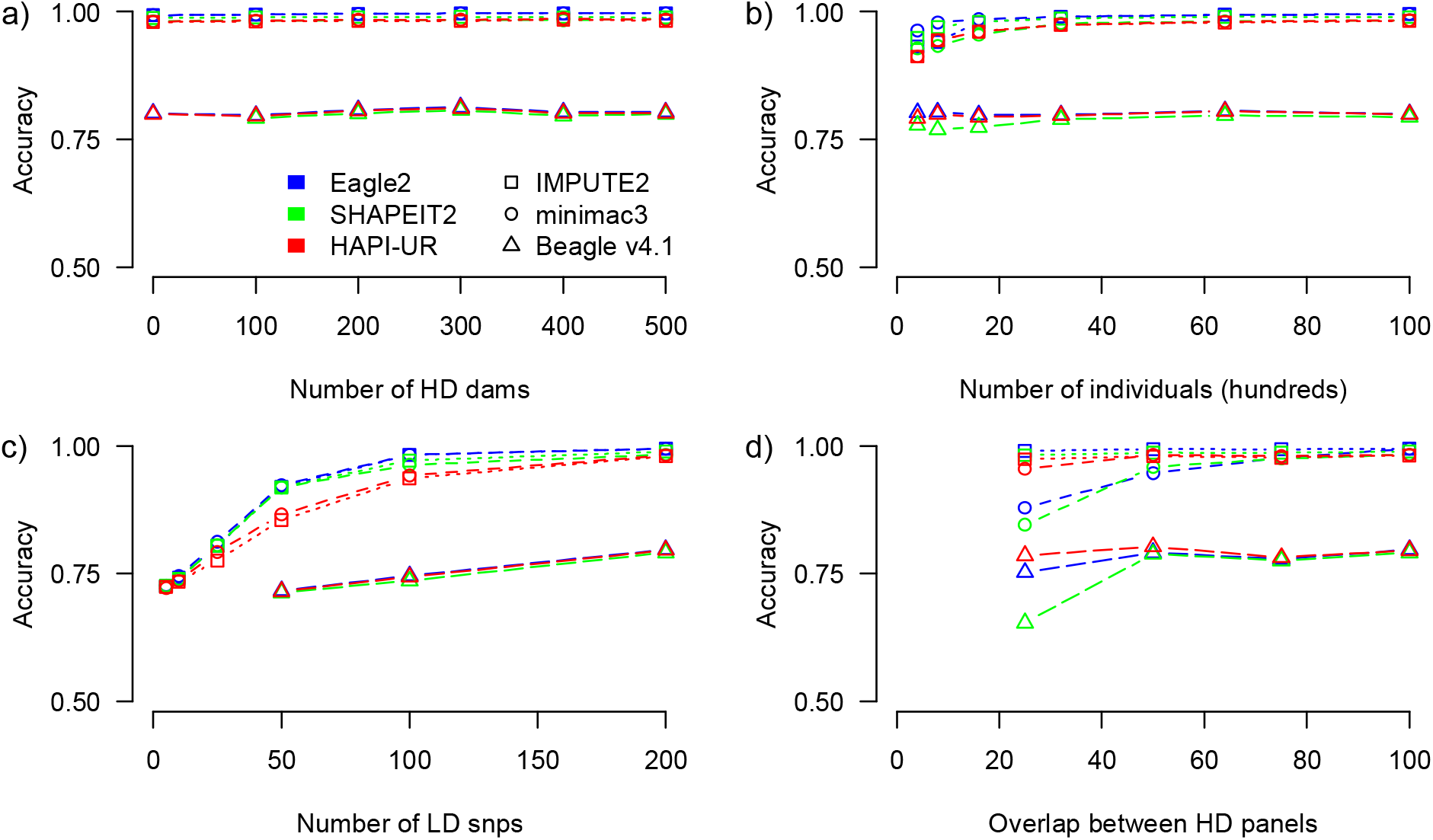
*Performance of each combination of pre-phasing and haploid HMM method. Unless otherwise noted there were 1000 high-density SNPs, 200 low-density SNPs, 100 dams genotyped at high-density and complete overlap between the high-density arrays of generations 1 and 2 and those of 3 and 4. We varied (a) the number of dams genotyped at high-density, (b) the number of individuals in the population, (c) the number of SNPs in the low-density array and (d) the amount of overlap between the high-density array for generations 1 and 2 and those of 3 and 4*.

### Run time and memory requirements

The elapsed run time of each method in the baseline scenario is given in Table 1. We found that of the diploid imputation methods, MaCH had the lowest run time followed by Beagle v4.0, fastPHASE, and IMPUTE2. Of the phasing methods, HAPI-UR was the fastest by an order of magnitude, followed by Eagle2 and SHAPEIT2. Of the haploid imputation methods, minimac3 was the fastest followed by Beagle v4.1 and IMPUTE2. The combined run-times of the two-step phasing and imputation methods were all substantially lower than that of the single step methods.

**Table 3.** Simulated data: Run time and accuracy for diploid imputation, phasing, and haploid imputation methods in the baseline scenario. The run time is given in seconds separately for phasing and imputation steps and as a total.

| Phasing method | Imputation method | HD Phasing (s) | LD Phasing (s) | Imputation (s) | Total (s) | Accuracy |
| --- | --- | --- | --- | --- | --- | --- |
| / | IMPUTE2 | / | / | 42,796 | 42,796 | 0.861 |
| / | Beagle v4.0 | / | / | 23,042 | 23,042 | 0.901 |
| / | MaCH | / | / | 21,998 | 21,998 | 0.944 |
| / | fastPHASE | / | / | 28,892 | 28,892 | 0.941 |
| HAPI-UR | IMPUTE2 | 117 | 14 | 149 | 280 | 0.964 |
| HAPI-UR | minimac3 | 117 | 14 | 62 | 193 | 0.967 |
| HAPI-UR | Beagle v4.1 | 117 | 14 | 78 | 209 | 0.793 |
| Eagle2 | IMPUTE2 | 1,361 | 207 | 148 | 1,717 | 0.988 |
| Eagle2 | minimac3 | 1,361 | 207 | 55 | 1,623 | 0.988 |
| Eagle2 | Beagle v4.1 | 1,361 | 207 | 79 | 1,647 | 0.794 |
| SHAPEIT2 | IMPUTE2 | 8,495 | 1,175 | 150 | 9,820 | 0.979 |
| SHAPEIT2 | minimac3 | 8,495 | 1,175 | 58 | 9,728 | 0.977 |
| SHAPEIT2 | Beagle v4.1 | 8,495 | 1,175 | 77 | 9,747 | 0.792 |

### Real Data

The performance on the real dataset was similar and is given in Table 4. The imputation accuracy of Eagle2 with minimac3 was 0.992, with Beagle v4.1 was 0.925, and with IMPUTE2 was 0.827. The imputation accuracy of HAPI-UR with minimac3 was 0.995%, with Beagle v4.1 was 0.939%, and with IMPUTE2 was 0.997%. Phasing with Eagle2 took 7 hours distributed across 8 cores. Phasing with HAPI-UR took 54 hours on a single core. All of the haploid imputation methods took under 6 hours. SHAPEIT2 was not able to phase the high-density and low-density individuals in 4 days and so was not analysed.

**Table 4.** Real data: Run time and accuracy for phasing, and haploid imputation methods on the real dataset scenario. The run time is given in hours separately for phasing and imputation steps and as a total. For Eagle2, the program was run distributed across 8 compute cores. HAPI-UR was run on a single core.

| Phasing method | Imputation method | HD Phasing (h) | LD Phasing (h) | Imputation (h) | Total (h) | Accuracy |
| --- | --- | --- | --- | --- | --- | --- |
| HAPI-UR | IMPUTE2 | 11.53 | 43.09 | 60.25 | 12.48 | 0.997 |
| HAPI-UR | minimac3 | 11.53 | 43.09 | 56.89 | 9.06 | 0.995 |
| HAPI-UR | Beagle v4.1 | 11.53 | 43.09 | 57.32 | 11.04 | 0.939 |
| Eagle2 | IMPUTE2 | 4.48 (8 cores) | 2.37 (8 cores) | 5.63 | 12.48 | 0.827 |
| Eagle2 | minimac3 | 4.48 (8 cores) | 2.37 (8 cores) | 2.21 | 9.06 | 0.992 |
| Eagle2 | Beagle v4.1 | 4.48 (8 cores) | 2.37 (8 cores) | 4.19 | 11.04 | 0.925 |

## Discussion

In this paper we evaluated the performance of HMM based imputation methods for imputation in animal populations. We found that combinations of phasing and haploid imputation methods provide increased imputation accuracy at substantially reduced runtimes compared to diploid imputation methods. The combination of using Eagle2 to pre-phase individuals and using minimac3 to impute the data lead to high accuracy imputation in a wide range of simulation scenarios and when analysing a real animal population.

The results of this paper highlight the power of separately phasing and imputing individuals. Intuitively it makes sense that performing phasing and imputation in a single step may increase imputation accuracy by marginalizing over uncertainty in phasing. However, the results here suggest that the additional accuracy lost by marginalizing over phasing errors is outweighed by the accuracy gained by considering larger haplotype reference panels. These results are particularly surprising in the context of animal populations where pre-existing reference panels may not exist (at least in the public domain), and so the reference panel itself is inferred by phasing high-density genotyped individuals. Our results suggest that modern phasing methods have a sufficiently high accuracy such that this phasing leads to only a small number of errors.

The performance of pre-phasing and haploid imputation is also surprising given the lower density of SNP arrays (both high-density and low-density), and the substantially lower number of overall individuals compared to human studies. We found that pre-phasing and haploid imputation was more effective than the best performing diploid imputation method even for a very small number of low-density markers or, low number of high-density dams, and low numbers of individuals.

Of the three phasing methods we tested, using Eagle2 led to the most accurate downstream imputation. This is likely due to the fact that Eagle2 is able to exploit longer segments of shared haplotypes between individuals, which are very common in highly related animal populations. Although Eagle2 led to the highest accuracy, we found that HAPI-UR was an order of magnitude faster for most datasets and resulted in a small decrease in accuracy on the simulated scenarios, but no decrease in accuracy on the real dataset. In their original paper, the authors of HAPI-UR suggest that it may be possible to increase the accuracy of HAPI-UR by running it multiple times with different window start positions and taking the consensus phase (Williams et al., 2012). Due to the low run time, this strategy would be feasible in animal populations but was not analysed here. SHAPEIT2, the oldest of the phasing methods had both the longest run-time which prevented us from evaluating it on the real dataset. Although the authors of SHAPEIT2 have now released SHAPEIT3, they do not recommend using it for populations of under 60,000 individuals and so the performance of SHAPEIT3 was not analysed here.

We found little difference in the performance of the assessed haploid imputation methods. Both Minimac3 and IMPUTE2 lead to accurate imputation. The accuracy of IMPUTE2 was consistently slightly (<1%) higher than that of minimac3 in simulated data, but the runtime was between two and three times that of minimac3. On the real dataset, the imputation accuracy of IMPUTE2 dropped when Eagle2 was used to pre-phase the data, but remained high when HAPI-UR was used to pre-phase the data. Overall the performance of Beagle v4.1 was poor for performing haploid imputation, although improved when analysing the real data set. This may be a result of the approximations used in Beagle v4.1, which were designed for imputation of human high-density SNP arrays to whole genome sequence data. These approximations seem less appropriate for low-density SNP arrays used in some animal populations.

With two exceptions, we found little interaction between the choice of phasing method and the choice of haploid imputation method. The first exception came in the performance of HAPI-UR when individuals were genotyped with multiple, semi-overlapping, SNP arrays. In this case the performance of HAPI-UR with minimac3 or Beagle v4.1 was substantially higher than the performance of Eagle2 with minimac3 or Beagle v4.1, although the accuracy of HAPI-UR with IMPUTE2 remained lower than that of Eagle2 with IMPUTE2. The underlying reason for this difference stems from the fact that in the case of minimac3 and Beagle v4.1 the phasing algorithms were also used to perform imputation on the missing non-overlapping SNPs in each high-density array, whereas in IMPUTE2 the two high-density arrays were phased separately, and IMPUTE2 was used to fill in missing SNPs as part of it’s high-density array merging step. The increased accuracy with HAPI-UR over Eagle2 in this scenario suggests that HAPI-UR can impute untyped loci in high-density arrays better than Eagle2. This is consistent with the second exception where HAPI-UR led to as high imputation accuracy, if not higher, as Eagle2 when performing imputation on the real dataset. Animals in the real dataset were genotyped with two high-density arrays, and two low-density arrays, and also exhibited a number of spontaneously missing SNPs. When using Eagle2 to phase individuals, IMPUTE2 and Beagle v4.1 markedly decreased in performance, particularly compared to minimac3. In contrast when HAPI-UR was used to phase individuals the performance of minimac3, IMPUTE2 and Beagle v4.1 remained high, suggesting an advantage of using HAPI-UR over Eagle2 when individuals are genotyped on multiple arrays or when observing a large amount of spontaneous missingness.

Some of the analysed phasing methods have an option to use pedigree information to improve phasing. Although these options were originally designed to help phase and impute parent-progeny trios (Browning and Browning, 2009), they can also be used for larger pedigrees (O’Connell et al., 2014). Previous work in phasing and imputing animal populations has found that combining pedigree and linkage information can improve phasing and imputation accuracy (Hickey et al., 2012). In this paper, we did not analyse the option to use pedigree information, but focused solely on HMMs based methods that use linkage-disequilibrium information for phasing and imputation as originally proposed by Li and Stephens (2003). SHAPEIT2 (O’Connell et al., 2014), Beagle v4.0 (Browning and Browning, 2009), and HAPI-UR (Williams et al., 2012) all provide options to use parent-progeny trio information. However, the two top performing methods, Eagle2 and minimac3, do not provide this option. Future work is needed to analyse how HMMs can utilize pedigree information to improve phasing and imputation, and to merge these insights with high-performance methods reviewed and tested here.

Overall, this study suggests that modern pre-phasing and haploid imputation methods can perform fast and accurate imputation of animal populations of any size. We noticed no disadvantage of using the two-step imputation approach even in cases of small populations, low-density SNP arrays, or multiple high-density arrays. Of the algorithms, we found that Eagle2 and HAPI-UR both reliably pre-phased the data and that IMPUTE2 and minimac3 lead to the highest imputation accuracy. However, we also noted a decreased accuracy when Eagle2 and IMPUTE2 were used to pre-phase and impute the data when animals were genotyped with semi-overlapping high-density SNP arrays. In this case the usage of Eagle 2 with minimac3 and HAPI-UR with IMPUTE2 or minimac3 lead to high accuracy. Overall, the results of these studies highlight the importance and feasibility of using HMMs to perform imputation in animal populations even as an increasing number of animals are genotyped and as genotyping densities increase.

## Acknowledgements

The authors acknowledge the financial support from the BBSRC ISPG to The Roslin Institute BB/J004235/1, from Genus PLC and from Grant Nos. BB/M009254/1, BB/L020726/1, BB/N004736/1, BB/N004728/1, BB/L020467/1, BB/N006178/1 and Medical Research Council (MRC) Grant No. MR/M000370/1. This work has made use of the resources provided by the Edinburgh Compute and Data Facility (ECDF) (http://www.ecdf.ed.ac.uk).

